# Optogenetic inhibition of the dorsal hippocampus CA3 region during early-stage cocaine-memory reconsolidation disrupts subsequent context-induced cocaine seeking in rats

**DOI:** 10.1101/2021.12.29.474477

**Authors:** Shuyi Qi, Shi Min Tan, Rong Wang, Jessica A. Higginbotham, Jobe L. Ritchie, Christopher K. Ibarra, Amy A. Arguello, Robert J. Christian, Rita A. Fuchs

## Abstract

The dorsal hippocampus (DH) is key to the long-term maintenance of cocaine memories following retrieval-induced memory destabilization; even though, it is not the site of protein synthesis-dependent memory reconsolidation. Here, we took advantage of the temporal and spatial specificity of an optogenetic manipulation to examine the role of the cornu ammonis 3 subregion of the DH (dCA3) in early-stage cocaine-memory reconsolidation. Male Sprague-Dawley rats expressing eNpHR3.0 in the DH were trained to self-administer cocaine in a distinct context and underwent extinction training in a different context. Rats then received a 15-min memory-reactivation session, to destabilize cocaine memories and trigger reconsolidation, or remained in their home cages (no-reactivation controls). Optogenetic inhibition of the dCA3 for 1 h immediately, but not 1 h, after memory reactivation resulted in cocaine-memory impairment as indicated by reduction in drug-seeking behavior selectively in the cocaine-paired context 3 d later, at test, relative to responding in no-inhibition, no-reactivation, and no-eNpHR3.0 controls. Cocaine-memory impairment was associated with reduced c-Fos expression, an index of neuronal activation, in the dCA3 stratum lucidum (SL) and stratum pyramidale (SP) at test. Based on these observations and extant literature, we postulate that recurrent circuits in the SP are activated during early-stage memory reconsolidation to maintain labile cocaine memories prior to protein synthesis-dependent restabilization in another brain region, such as the basolateral amygdala. Furthermore, SL and SP interneurons may enhance memory reconsolidation by limiting synaptic noise in the SP and also contribute to recall as elements of the updated cocaine engram or retrieval links.

## INTRODUCTION

Cocaine use disorder is characterized by uncontrollable drug craving and high relapse propensity^1,2^. In the course of chronic drug use, associations are formed between distinct environmental contexts, responses that lead to drug procurement, and the unconditioned motivational effects of cocaine^3^. These context-response-cocaine associations are consolidated into long-term memory^4^. Thus, exposure to environmental contexts that predict drug availability (i.e., cocaine-paired contexts) can lead to the retrieval of cocaine memories and to robust incentive motivation to seek cocaine^1,5^. Importantly, retrieval can also result in the destabilization of cocaine memories^4^. Destabilized memories are susceptible to manipulation^6^, and their maintenance requires reconsolidation into long-term memory stores through a process that involves *de novo* protein synthesis and glutamatergic synaptic plasticity^7^. Empirical evidence indicates that interference with drug-memory reconsolidation transiently reduces craving in human subjects^8,9^ and drug-seeking behavior in animal models of drug relapse^10-12^. Therefore, it is of interest from an anti-relapse treatment perspective to understand the neural substrates recruited for cocaine-memory reconsolidation.

The dorsal hippocampus (**DH**) is a brain region involved in the reconsolidation of spatial, contextual-fear, and contextual-drug memories^13-16^. Inhibition of mRNA synthesis in the DH during spatial-memory reconsolidation disrupts memory integrity in rats, as reflected by impaired Morris water maze performance^14^. Similarly, zif268 knockdown in the DH during fear-memory reconsolidation attenuates conditioned fear-memory strength, as indicated by reduced freezing behavior^13^. We have shown that inhibition of neural conductance or Src-family tyrosine kinase activity in the DH after cocaine-memory reactivation diminishes, whereas anisomycin-induced inhibition of *de novo* protein synthesis fails to alter, cocaine-memory strength as indicated by attenuated drug context-induced cocaine-seeking behavior^16-17^. These observations suggest that the DH is not the site of memory restabilization in our model. Instead, the DH may support protein synthesis-dependent memory restabilization in other brain regions during reconsolidation.

Associative microcircuits in the cornu ammonis 3 subregion of the DH (**dCA3**) have been theorized to be critical for short-term memory retrieval, memory consolidation, long-term memory recall^18,19^, and, more recently, memory reconsolidation^20^. Consistent with this, the dCA3 exhibits cAMP response element-binding protein activation at contextual fear-memory recall, and this response is inhibited by treatments that interfere with fear-memory reconsolidation^21^. We have hypothesized that the dCA3 maintains destabilized cocaine memories and, therefore, it is recruited transiently, during the early stages of memory reconsolidation, when most memory traces are still labile. We took advantage of the temporal and spatial precision of optogenetics to inhibit neuronal activity in the dCA3 during early-or later-stage cellular memory reconsolidation, which we operationally defined as the first and second h of memory processing after cocaine-memory reactivation, respectively. We examined the resulting effects on long-term memory strength as indicated by drug context-induced cocaine-seeking behavior three days later. We predicted that optogenetic inhibition of the dCA3 during early-stage memory reconsolidation would be sufficient to diminish cocaine-memory strength. Furthermore, we explored the possible contributions of the stratum lucidum (**SL**) and stratum pyramidale (**SP**) cell layers of the dCA3 to this phenomenon.

## MATERIALS AND METHODS

### Animals

Male Sprague-Dawley rats (N= 74; 275–300 g; Envigo, South Kent, WA) were maintained in a climate-controlled vivarium on a reversed light-dark cycle. The rats received water *ad libitum* and 20–25 g of rat chow per day. Animal-housing and treatment protocols followed the *Guide for the Care and Use of Laboratory Rats* (Institute of Laboratory Animal Resources on Life Sciences, 2011) and were approved by the Washington State University Institutional Animal Care and Use Committee.

### Surgery

Rats were anesthetized using ketamine and xylazine (100.0 mg/kg and 5.0 mg/kg, i.p., respectively) 24 h after a food-training session (see **Supplemental Methods**). AAV5-hSyn-eNpHR3.0-eYFP-WPRE-PA (5.5×10e12 vm/ml; University of North Carolina Vector Core, Chapel Hill, NC) or AAV5-hSyn-eYFP (3.5×10e12 vm/ml; University of North Carolina Vector Core) was infused bilaterally into the DH (−3.4 mm AP, +/-3.97 mm ML, −3.5 mm DV from bregma; 15°) at a volume of 0.7 μl/hemisphere over 10 min. Next, optic fibers (200-μm diameter fiber optic cable incased in a 6.4-mm long ceramic ferrule; Thor Labs, Newton, NJ) were aimed at the dCA3 (−3.4 mm AP, +/-3.1 mm ML, −3.58 mm DV; 0°). Stainless-steel screws and dental acrylic secured the optic fibers to the skull. Rats underwent jugular catheter implantation surgery up to 3 days later, as described previously^22^. Virus microinfusion and catheter surgery protocols are described in **Supplemental Methods**. Rats received a bacon-flavored placebo tablet (5g tablet; Bio-Serv) 24 h prior to the first surgery and bacon-flavored Rimadyl® MD tablets (2 mg carprofen/5g tablet/day; Bio-Serv) for at least 48 h after each surgery.

### Drug Self-Administration and Extinction Training

Rats were randomly assigned to receive 2-h cocaine self-administration training sessions in one of two environmental contexts (context 1 or context 2, described in **Supplemental Methods**; see experimental timelines in **Fig. 1-5A**). Rats were trained to press an “active” lever under a fixed-ratio 1 cocaine-reinforcement schedule (cocaine hydrochloride, 0.5 mg/ml, 50 μl/infusion; NIDA Drug Supply Program, Research Triangle Park, NC) with a 20-sec timeout period. Lever presses on a second, “inactive,” lever and all lever presses during the timeout period had no programmed consequences. Training sessions continued daily until the rats obtained ≥ 10 infusions/session on 10 days. Rats then receive seven extinction training sessions (2 h/day) in the context (1 or 2) that had not been used for cocaine self-administration training. During extinction training, lever presses had no programmed consequences. After extinction sessions 5-7, rats were acclimated to the optogenetic procedure, as described in **Supplemental Methods**.

**Figure 1.**
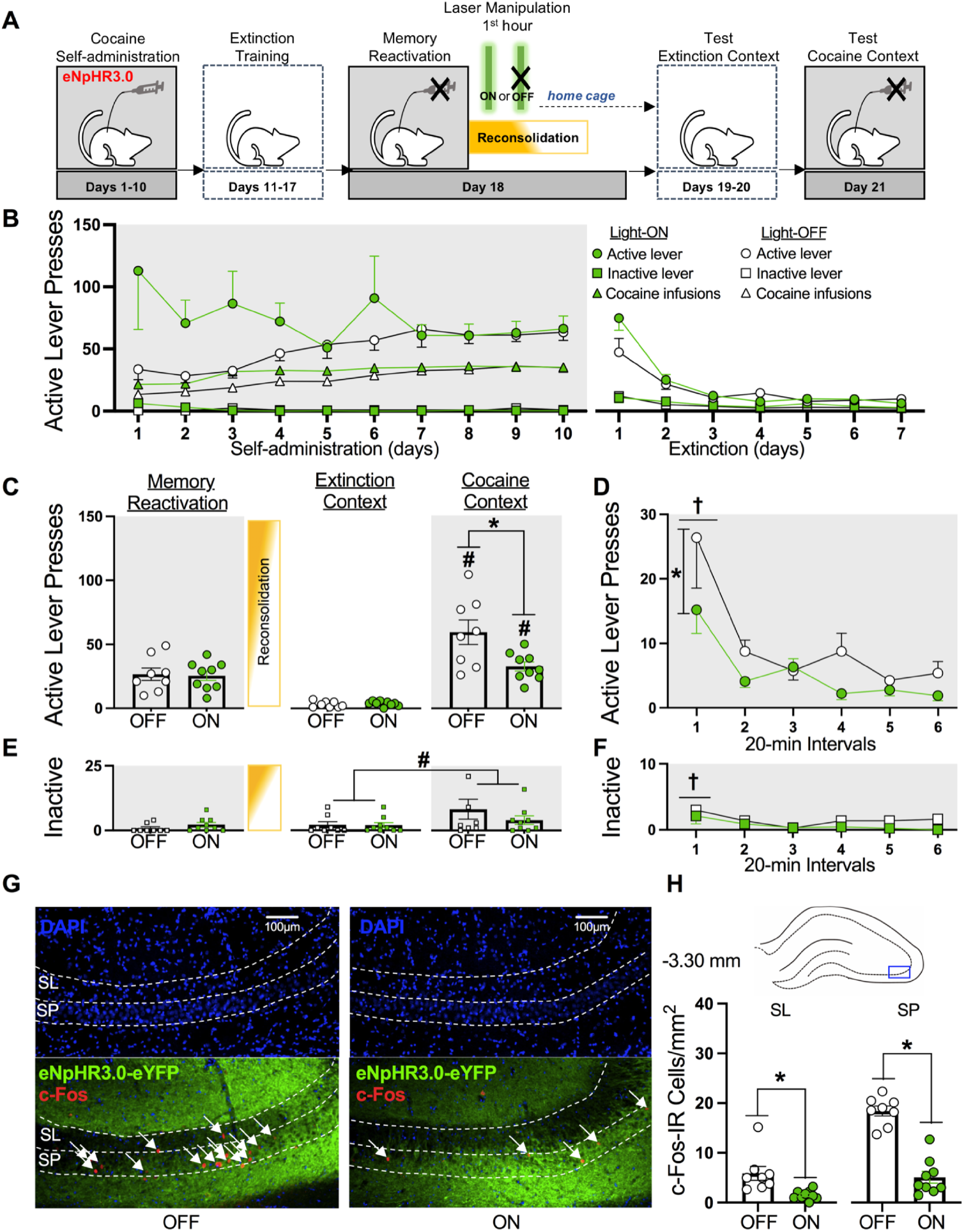
Optogenetic inhibition of the dCA3 during the first h of cocaine-memory reconsolidation reduces cocaine-memory strength and dCA3 neuronal activation at test. (**A**) Experimental timeline. Following cocaine self-administration and extinction training, eNpHR3.0-expressing rats received a 15-min memory-reactivation session, immediately followed by Laser-ON (n = 9) or Laser-OFF (n = 8) treatment for 1 h. Lastly, rats received test sessions in the extinction and cocaine-paired contexts to assess extinction- and cocaine-memory strength, respectively. (**B**) Active- and inactive-lever presses and cocaine infusions (mean/2 h ± SEM) by the Light-ON and Light-OFF groups during cocaine self-administration training (last 10 days) and extinction training. (**C**) Active-lever presses during the memory-reactivation session (mean/15 min ± SEM) and upon first post-treatment re-exposure to the extinction and cocaine-paired contexts at test (mean/2 h ± SEM). Symbols: ANOVA ^**#**^context simple-main effect, *p* < 0.001; **\***treatment simple-main effect, *p* = 0.02. (**D**) Time-course of active-lever presses at test in the cocaine-paired context (mean/20-min interval ± SEM). Symbols: ANOVA ^**†**^time main effect, Tukey’s tests, interval 1 > intervals 2-6, *p*s < 0.05; **\***treatment main effect, *p* = 0.02. (**E**) Inactive-lever presses (mean ± SEM) during the memory-reactivation session (mean/15 min ± SEM) and upon first post-treatment re-exposure to the extinction and cocaine-paired contexts at test (mean/2 h ± SEM). Symbol: ANOVA ^**#**^context main effect, *p* = 0.03. (**F**) Time-course of inactive-lever presses at test in the cocaine-paired context (mean/20-min interval ± SEM). Symbol: ANOVA ^**†**^time main effect, Tukey’s tests, interval 1 > intervals 2-6, *p*s < 0.05. (**G**) Representative 20x photomicrographs of the dCA3 stratum lucidum (SL) and stratum pyramidale (SP) of rats in the Light-OFF and Light-ON groups. Brain tissue was collected immediately after the test in the cocaine-paired context. Images showing DAPI staining (blue) were used to visualize SL and SP boundaries (see also Fig. S1), and corresponding overlay images were used to visualize eNpHR3.0-eYFP expression (green) and c-Fos immunoreactive (IR) cell bodies (red, *arrows*). (**H**) The density of c-Fos-immunoreactive (IR) neurons was quantified in the area indicated by the blue rectangle on the brain schematic. c-Fos-IR cell body density (mean ± SEM) in the SL and SP of rats in the Light-ON and Light-OFF groups. Symbols: **\****t*-tests, *p*s < 0.05.

### Memory Reactivation

Twenty-four h after the last extinction-training session, all rats in experiments 1, 3, 4, and 5 were exposed to the previously cocaine-paired context for 15 min to destabilize cocaine memories and trigger reconsolidation. During the session, lever presses had no programmed consequences. All rats in experiment 2 remained in their home cages during this time (no-memory reactivation).

### In vivo Optogenetics

Immediately or 1 h after memory reactivation or no-memory reactivation, all rats were placed into black metal chambers to which they had been acclimated. Their optic fibers were connected to 532-nM solid-state lasers (Shanghai Laser & Optics Century Company, Shanghai, China) through fiber-optic patch cables (Thor Labs) and an optical commutator (Doric Lenses, Quebec, CAN). Rats received Light-ON or Light-OFF treatment (n = 7-9/group) with treatment assignment balanced based on mean active-lever responding and cocaine intake during the last three drug self-administration sessions. Light-ON treatment consisted of laser-light stimulation (10 mW, 5 sec on, 1 sec off) for 1 h, whereas Light-OFF treatment involved no laser-light stimulation. Rats in experiment 5 were euthanized immediately after optogenetic treatment.

### Testing in the Extinction and Cocaine-paired Contexts

Twenty-four h after optogenetic treatment, daily 2-h extinction-training sessions resumed in experiment 1-4 and continued until active-lever responding declined to ≤ 25 responses/session on two consecutive sessions. Twenty-four h later, the rats were placed into the previously cocaine-paired context for a 2-h session. During test sessions, lever presses had no programmed consequences. Lever responses in the extinction context (first session post treatment) and cocaine-paired context were used to index treatment effects on extinction-memory strength (off target) and cocaine-memory strength, respectively.

### Brain Histology and Immunohistochemistry

Rats were overdosed with ketamine and xylazine (300/15 mg/kg, i.p., respectively) then transcardially perfused with 1X ice-cold phosphate-buffered saline (PBS, 100 ml) then 4% formaldehyde (100 ml; Sigma). Brains were dissected out, fixed using 4% formaldehyde at 4 C° for 24 h, and cryoprotected with 30% sucrose/0.1% sodium azide solution at 4 C°. The brains were stored at − 80 C° until they were cut into 30-μm coronal sections.

In experiments 1-4, brain tissue was collected immediately after the last 2-h test session to visualize virus expression, optic-fiber placement, and c-Fos expression in the SL and SP of the dCA3 at test. dCA3 cell layers were visualized using intensity differences in DAPI staining^23^ (**Fig. S1F**). In experiment 5, brain tissue was collected immediately after memory reactivation plus the 1-h optogenetic manipulation to verify that Light-ON treatment reduced c-Fos expression in the dCA3 during memory reconsolidation and to investigate the cell types affected. Standard immunohistochemistry and microscopy methods and antibody information are described in **Supplemental Methods**.

### Data Analysis

Potential pre-existing group differences in drug infusions and lever presses during the training sessions were analyzed using mixed-factorial analyses of variance (ANOVAs) with treatment (Light-ON, Light-OFF) as between-subjects factor and time (day) as within-subject factor. Potential pre-existing group differences in lever presses during the memory-reactivation session were analyzed using independent-samples *t*-tests. Treatment effects on lever presses upon first re-exposure to the extinction and cocaine-paired contexts at test were assessed using mixed-factorial ANOVAs with treatment as between-subjects factor and context (cocaine-paired, extinction) and time (20-min intervals) as within-subject factors, where appropriate. Significant interaction and time main effects were further investigated using *post hoc* Tukey’s or Bonferroni tests (adjusted *α* = 0.01). Treatment effects on c-Fos expression were examined using independent-samples *t*-tests. Alpha was set at 0.05.

## RESULTS

### Histology and Behavioral History

eNpHR3.0-eYFP and eYFP expression were observed in the dCA1-3 in experiments 1-5 (**Fig. S1A-E**). Optic-fiber tracts were located in the dCA3 for all rats whose data were included in data analysis. Eighteen rats were excluded based on optic-fiber misplacement or unilateral/insufficient virus expression.

There were no statistically significant pre-existing group differences in cocaine intake or in active- or inactive-lever responding during cocaine self-administration and extinction training (**Figs. 1B-4B**), or during the cocaine-memory reactivation session (**Figs. 1C, 3C, 4C**) (**Table S1**). Furthermore, optogenetic treatment did not alter the number of extinction test sessions (mean ± SEM = 2.30 ± 0.07 sessions; **Fig. 1C-4C**).

### Optogenetic Inhibition of the dCA3 during the First h after Memory Reactivation Reduces Cocaine-Memory Strength

Experiment 1 examined whether neuronal activity in the dCA3 during the first h of cocaine-memory reconsolidation was necessary for cocaine-memory maintenance, as indexed by the magnitude of drug-seeking behavior in the cocaine-paired context at test (**Fig. 1**). Optogenetic dCA3 inhibition during the first h after cocaine-memory reactivation reduced active-lever responding at test in a context-specific manner (**Fig. 1C**; 2 × 2 ANOVA, context x treatment interaction *F*_(1,15)_ = 8.22, *p* = 0.01; treatment main *F*_(1,15)_ = 6.75, *p* = 0.02; context main *F*_(1,15)_ = 79.20, *p* < 0.001). Specifically, three days after Light-ON (Bonferroni *t*_(8)_ = 7.0, *p* < 0.001) or Light-OFF (Bonferroni *t*_(7)_ = 6.21, *p* < 0.001) treatment, cocaine-paired context exposure significantly increased active-lever responding at test compared to extinction-context exposure. Furthermore, Light-ON treatment reduced active-lever responding in the cocaine-paired context (Bonferroni *t*_(15)_ = −2.74, *p* = 0.02), but not in the extinction context (Bonferroni *t*_(15)_ = 0.64, *p* = 0.53), compared to Light-OFF treatment. Active-lever responding decreased over time in the cocaine-paired context at test, independent of treatment, and Light-ON treatment reduced active-lever responding compared to Light-OFF treatment, independent of time (**Fig. 1D**; 2 × 6 ANOVA, time main *F*_(5,75)_ = 12.01, *p* < 0.001, Tukey’s tests, interval 1 > intervals 2-6, *p*s < 0.05; treatment main *F*_(1,15)_ = 7.52, *p* = 0.02; treatment x time interaction *F*_(5,75)_ = 1.19, *p* = 0.32).

Cocaine-paired context exposure increased inactive-lever responding at test relative to extinction-context exposure independent of treatment (**Fig. 1E**; ANOVA, context main *F*_(1,15)_ = 5.84, *p* = 0.03; treatment main and treatment x context interaction *F*s_(1,15)_ ≤ 1.76, *p*s ≥ 0.20). Furthermore, inactive-lever responding decreased over time in the cocaine-paired context at test independent of treatment (**Fig. 1F**; ANOVA, time main *F*(_5,75)_ = 2.42, *p* = 0.04, Tukey’s tests, interval 1 > interval 3, *p*s < 0.05; treatment main and treatment x time interaction *F*s_(1-5,12-75)_ ≤ 1.62, *p*s ≥ 0.22).

In brain tissue collected immediately after the 2-h test session in the cocaine-paired context, c-Fos expression in the dCA3 SL and SP was quantified ventral to the optic fiber tract. Light-ON treatment during the first h after cocaine-memory reactivation reduced c-Fos expression in the SL (*t*_(15)_ = 3.20, *p* < 0.01) and SP (*t*_(15)_ = 8.45, *p* < 0.001) at test, compared to Light-OFF treatment (**Fig. 1G-H**).

### Optogenetic Inhibition of the dCA3 Without Memory Reactivation does not Alter Cocaine-Memory Strength

Experiment 2 examined whether effects in experiment 1 reflected a genuine, memory-reactivation-dependent^24^ reconsolidation deficit by evaluating whether optogenetic dCA3 inhibition without cocaine-memory reactivation would impair cocaine-memory strength (**Fig. 2**). Cocaine-paired context exposure increased active-lever responding relative to extinction-context exposure, and Light-ON treatment administered for 1 h immediately after home cage stay (i.e., no-memory reactivation) did not alter active-lever responding in either context, relative to Light-OFF treatment (**Fig. 2C**; ANOVA, context main *F*_(1,12)_ = 35.32, *p* < 0.001; treatment main and treatment x context interaction *F*s_(1,12)_ ≤ 0.81, *p*s ≥ 0.39). Active-lever responding decreased over time in the cocaine-paired context at test, independent of treatment (**Fig. 2D**; ANOVA, time main *F*_(5,60)_ = 24.83, *p* < 0.0001, Tukey’s tests, interval 1 > intervals 2-6, *p*s < 0.05; treatment main and treatment x time interaction *F*s_(1-5,12-60)_ ≤ 0.86, *p*s ≥ 0.49).

**Figure 2.**
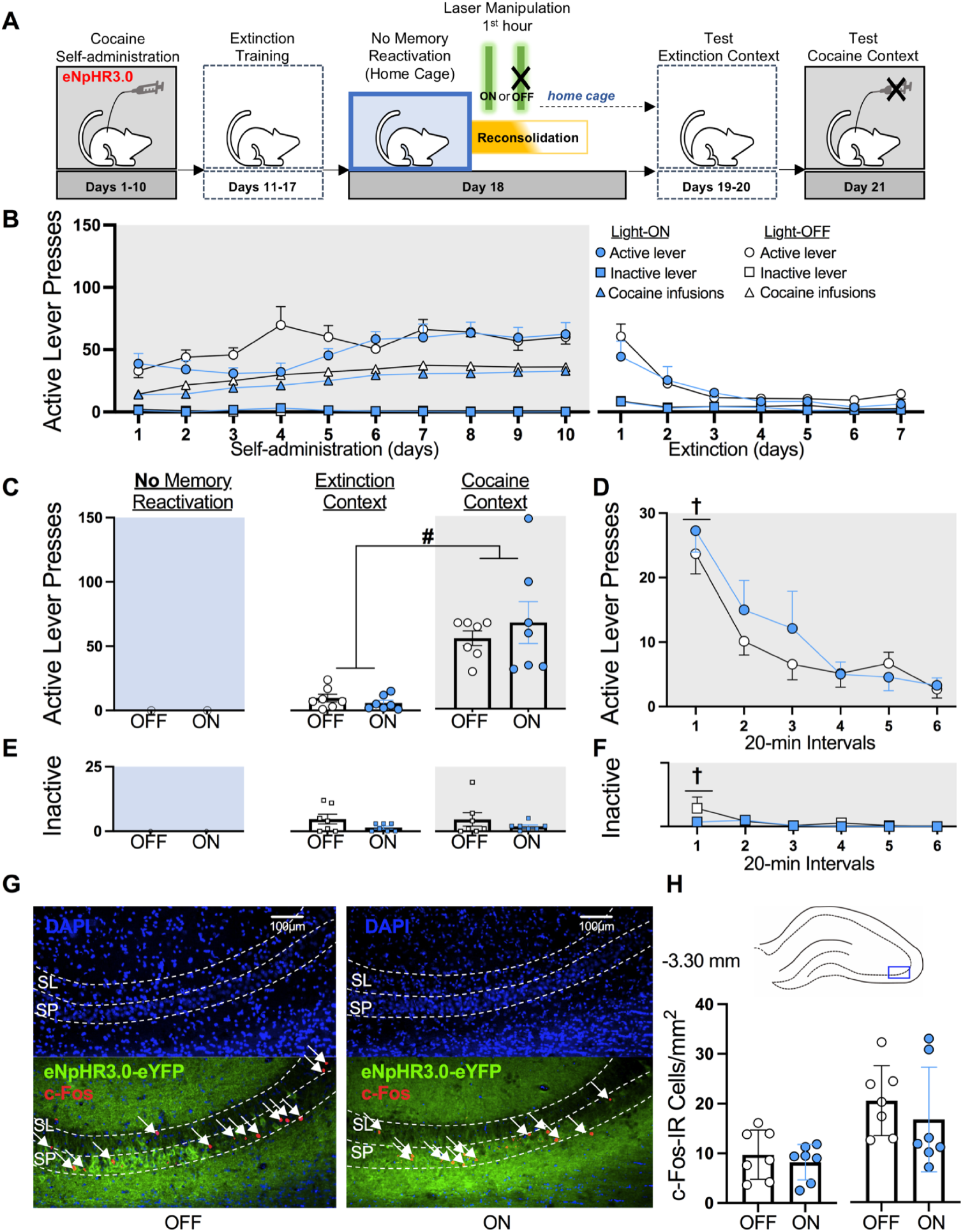
Optogenetic inhibition of the dCA3 without memory reactivation fails to alter cocaine-memory strength or dCA3 neuronal activation at test. (**A**) Experimental timeline. Following cocaine self-administration and extinction training, eNpHR3.0-expressing rats remained in their home cages (i.e., no memory reactivation), then received Laser-ON (n = 7) or Laser-OFF (n = 7) treatment for 1 h. Lastly, rats received test sessions in the extinction and cocaine-paired contexts to assess extinction- and cocaine-memory strength, respectively. (**B**) Active- and inactive-lever presses and cocaine infusions (mean/2 h ± SEM) by the Light-ON and Light-OFF groups during cocaine self-administration training (last 10 days) and extinction training. (**C**) Active-lever presses upon first post-treatment re-exposure to the extinction and cocaine-paired contexts at test (mean/2 h ± SEM). Symbol: ANOVA ^**#**^context main effect, *p* < 0.001. (**D**) Time-course of active-lever presses in the cocaine-paired context at test (mean/20-min interval ± SEM). Symbol: ANOVA ^**†**^time main effect, Tukey’s tests, interval 1 > intervals 2-6, *p*s < 0.05. (**E**) Inactive-lever presses (mean ± SEM) upon first post-treatment re-exposure to the extinction and cocaine-paired contexts at test (mean/2 h ± SEM). (**F**) Time-course of inactive-lever presses in the cocaine-paired context at test (mean/20-min interval ± SEM). Symbol: ANOVA ^**†**^time main effect, intervals 1> intervals 4-6, *p*s < 0.05. (**G**) Representative 20x photomicrographs of the dCA3 stratum lucidum (SL) and stratum pyramidale (SP) of rats in the Light-OFF and Light-ON groups. Brain tissue was collected immediately after the test session in the cocaine-paired context. Images showing DAPI staining (blue) were used to visualize SL and SP boundaries (see also Fig. S1), and corresponding overlay images were used to visualize eNpHR3.0-eYFP expression (green) and c-Fos immunoreactive (IR) cell bodies (red, *arrows*). (**H**) The density of c-Fos-immunoreactive (IR) neurons was quantified in the area indicated by the blue rectangle on the brain schematic. c-Fos-IR cell body density (mean ± SEM) in the SL and SP of rats in the Light-ON and Light-OFF groups.

Inactive-lever responding did not vary by testing context or treatment (**Fig. 2E**; ANOVA, all *F*s_(1,12)_ ≤ 1.94, *p*s ≥ 0.76). Inactive-lever responding decreased across time in the cocaine-paired context at test, independent of treatment (**Fig. 2F**; 2 × 6 ANOVA, time main *F*_(5,60)_ = 2.88, *p* = 0.02, Tukey’s tests, interval 1 > intervals 4-6, *p*s < 0.05; treatment main and treatment x time interaction *F*s_(1-5,12-60)_ ≤ 1.10, *p*s ≥ 0.33).

Light-ON treatment administered for 1 h after no-memory reactivation did not alter c-Fos expression in the SP (*t*_(12)_ = 0.79, *p* = 0.44) or SL (*t*_(12)_ = 0.65, *p* = 0.53) at test, compared to Light-OFF treatment (**Fig. 2G-H**).

### Optogenetic Inhibition of the dCA3 during the Second h after Memory Reactivation Fails to Alter Cocaine-Memory Strength

Experiment 3 examined whether optogenetic dCA3 inhibition during the second h of memory reconsolidation would weaken cocaine-memory strength (**Fig. 3**). Cocaine-paired context exposure increased active-lever responding relative to extinction-context exposure, and Light-ON treatment during the second h after memory reactivation failed to alter active-lever responding in either context relative to Light-OFF treatment (**Fig. 3C**; ANOVA, context main *F*_(1,14)_ = 40.78, *p* < 0.001; treatment main and treatment x context interaction *F*s_(1,14)_ ≤ 0.87, *p*s ≥ 0.37). Furthermore, active-lever responding decreased over time in the cocaine-paired context, at test, independent of treatment (**Fig. 3D**; ANOVA, time main *F*_(5,70)_ = 13.10, *p* < 0.001, Tukey’s test, interval 1 > intervals 3-6, *p*s < 0.05; treatment main and treatment x time interaction *F*s_(1-5,14-70)_ ≤ 0.19, *p*s ≥ 0.67).

**Figure 3.**
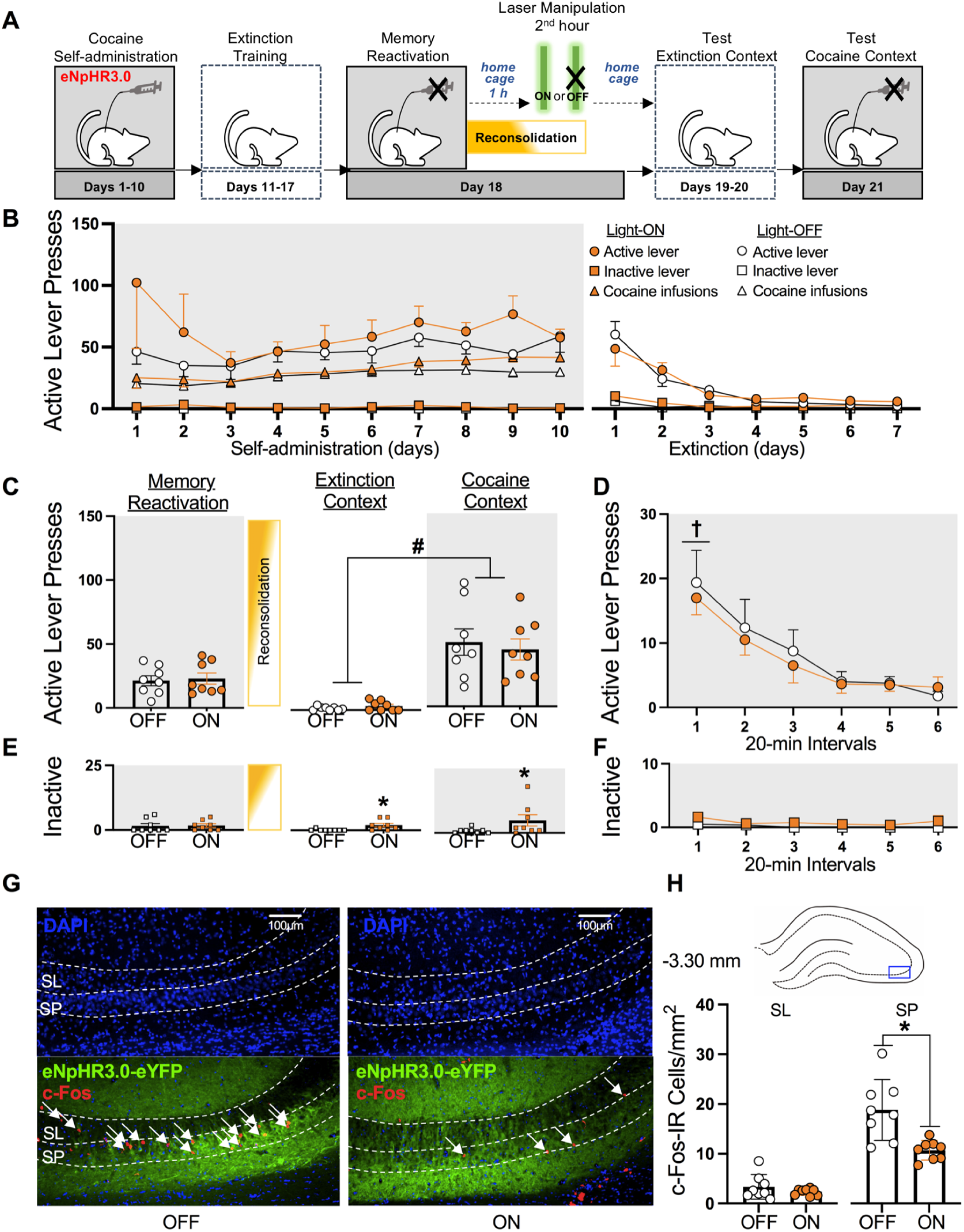
Optogenetic inhibition of the dCA3 during the second h of memory reconsolidation does not alter cocaine memory strength and reduces dCA3 neuronal activity in a region-specific manner at test. (**A**) Experimental timeline. Following cocaine self-administration and extinction training, eNpHR3.0-expressing rats received a 15-min memory-reactivation session. Rats were returned to their home cages for 1 h and then received Laser-ON (n = 8) or Laser-OFF (n = 8) treatment for 1 h. Lastly, rats received test sessions in the extinction and cocaine-paired contexts to assess extinction- and cocaine-memory strength, respectively. (**B**) Active- and inactive-lever presses and cocaine infusions (mean/2 h ± SEM) by the Light-ON and Light-OFF groups during cocaine self-administration training (last 10 days) and extinction training. (**C**) Active-lever presses during the memory-reactivation session (mean/15 min ± SEM) and upon first post-treatment re-exposure to the extinction and cocaine-paired contexts at test (mean/2 h ± SEM). Symbol: ANOVA ^**#**^context main effect, *p* < 0.001. (**D**) Time-course of active-lever presses in the cocaine-paired context at test (mean/20-min interval ± SEM). Symbol: ANOVA ^**†**^time main effect, Tukey’s test, interval 1 > intervals 3-6, *p*s < 0.05; **\***treatment main effect, *p* = 0.05. (**E**) Inactive-lever presses (mean ± SEM) during the memory-reactivation session (mean/15 min ± SEM) and upon first post-treatment re-exposure to the extinction and cocaine-paired contexts at test (mean/2 h ± SEM). Symbols: ANOVA **\***treatment main effect, *p* < 0.01. (**F**) Time-course of inactive-lever presses in the cocaine-paired context at test (mean/20-min interval ± SEM). (**G**) Representative 20x photomicrographs of the dCA3 stratum lucidum (SL) and stratum pyramidale (SP) of rats in the Light-OFF and Light-ON groups. Brain tissue was collected immediately after the test session in the cocaine-paired context. Images showing DAPI staining (blue) were used to visualize SL and SP boundaries (see also Fig. S1), and corresponding overlay images were used to visualize eNpHR3.0-eYFP expression (green) and c-Fos immunoreactive (IR) cell bodies (red, *arrows*). (**H**) The density of c-Fos-immunoreactive (IR) neurons was quantified in the area indicated by the blue rectangle on the brain schematic. c-Fos-IR cell body density (mean ± SEM) in the SL and SP of rats in the Light-ON and Light-OFF groups. Symbol: **\****t*-test, *p*s < 0.05.

Light-ON treatment in the dCA3 during the second h after memory reactivation increased inactive-lever responding compared to Light-OFF treatment, independent of testing context (**Fig. 3E**; ANOVA, treatment main *F*_(1,14)_ = 9.78, *p* < 0.01; context main and treatment x context interaction *F*s_(1,14)_ ≤ 1.34, *p*s ≥ 0.27). However, inactive-lever responding in the cocaine-paired context at test did not vary as a function of treatment type or time (**Fig. 3F**; ANOVA, all *F*s_(1-5,14-70)_ ≤ 3.35, *p*s ≥ 0.09).

Light-ON treatment administered during the second h after memory reactivation reduced c-Fos expression in the SP (*t*_(14)_ = 7.62, *p* < 0.001), but not in the SL (*t*_(14)_ = 1.94, *p* = 0.07), at test, compared to Light-OFF treatment (**Fig. 3G-H**).

### Laser-ON Treatment Without eNpHR3.0 Expression in the dCA3 does not Alter Cocaine-Memory Strength

Experiment 4 evaluated whether Light-ON treatment in eYFP-expressing rats (i.e., without eNpHR3.0) during the first h of memory reconsolidation would impair cocaine-memory strength (**Fig. 4**) to examine whether the effects observed in experiment 1 reflected laser light-induced nonspecific performance deficit (e.g., heat-induced cell damage^25^). Cocaine-paired context exposure increased active-lever responding relative to extinction-context exposure at test, independent of treatment (**Fig. 4C**; ANOVA, context main *F*_(1,14)_ = 60.95, *p* < 0.001; treatment main and treatment x context interaction *F*s_(1,14)_ ≤ 0.51, *p*s ≥ 0.49). Thus, Light-ON treatment without eNpHR3.0 did not alter active-lever responding in either context relative to Light-OFF treatment. Active-lever responding decreased over time in the cocaine-paired context at test, independent of treatment (**Fig. 4D**; ANOVA, time main *F*_(5,70)_ = 18.99, *p* < 0.001, Tukey’s test interval 1 > intervals 2-6, *p*s < 0.05; treatment main and treatment x time interaction *F*s_(1-5,14-70)_ ≤ 0.34, *p*s ≥ 0.80).

**Figure 4.**
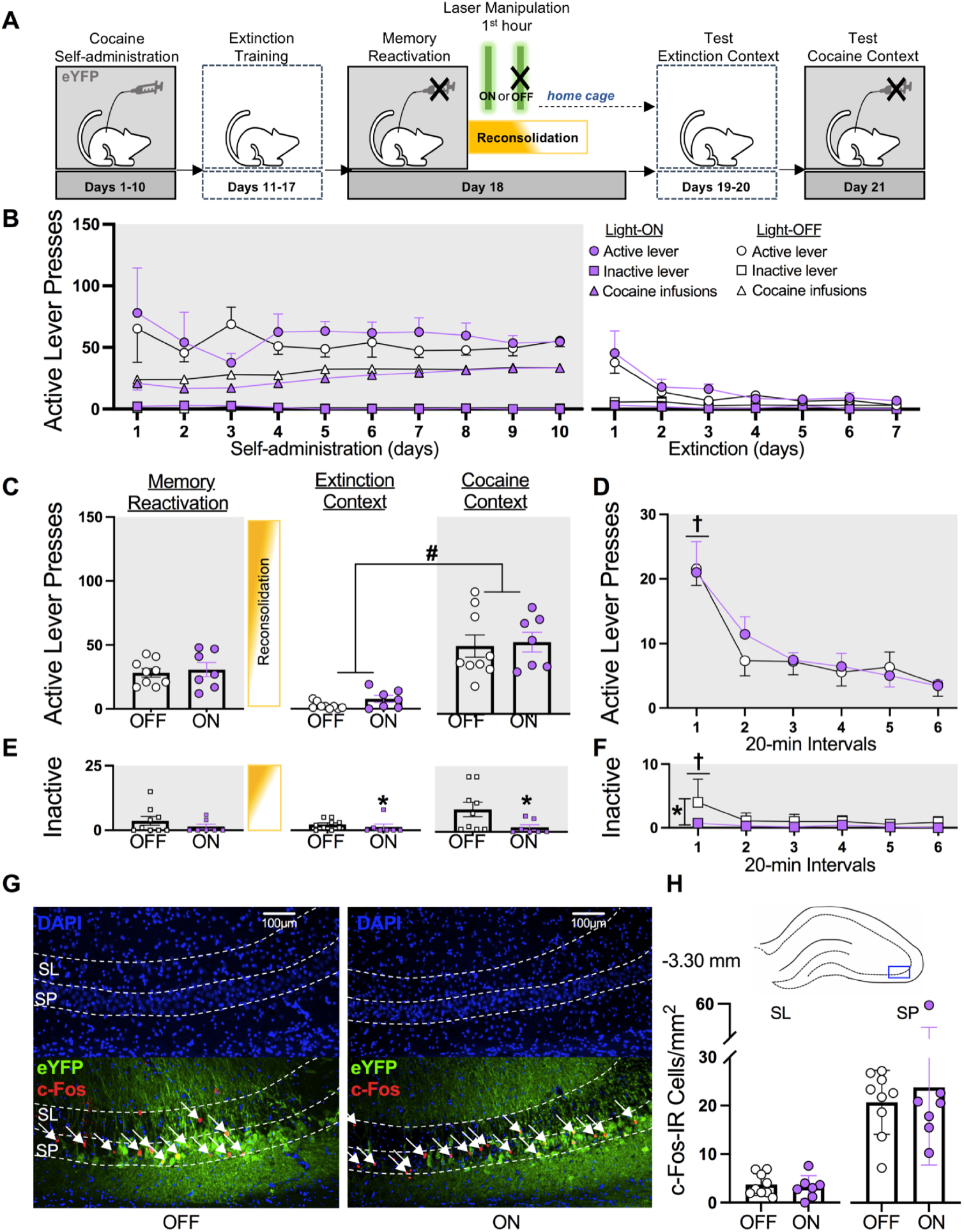
Light-ON treatment without eNpHR3.0 expression in the dCA3 does not alter cocaine-memory strength or dCA3 neuronal activity at test. (**A**) Experimental timeline. Following cocaine self-administration and extinction training, eYFP-expressing rats received a 15-min memory-reactivation session, immediately followed by Laser-ON (n = 7) or Laser-OFF (n = 9) treatment for 1 h. Lastly, rats received test sessions in the extinction and cocaine-paired contexts to assess extinction- and cocaine-memory strength, respectively. (**B**) Active- and inactive-lever presses and cocaine infusions (mean/2 h ± SEM) by the Light-ON and Light-OFF groups during cocaine self-administration training (last 10 days) and extinction training. (**C**) Active-lever presses during the memory-reactivation session (mean/15 min ± SEM) and upon first post-treatment re-exposure to the extinction and cocaine-paired contexts at test (mean/2 h ± SEM). Symbols: ANOVA ^**#**^context main effect, *p* < 0.001. (**D**) Time-course of active-lever presses in the cocaine-paired context at test (mean/20-min interval ± SEM). Symbols: ANOVA ^**†**^time main effect, Tukey’s test interval 1 > intervals 2-6, *p*s < 0.05. (**E**) Inactive-lever presses (mean ± SEM) during the memory-reactivation session (mean/15 min ± SEM) and upon first post-treatment re-exposure to the extinction and cocaine-paired contexts at test (mean/2 h ± SEM). Symbol: ANOVA ^**#**^treatment main effect, *p* = 0.03. (**F**) Time-course of inactive-lever presses in the cocaine-paired context at test (mean/20-min interval ± SEM). Symbol: ANOVA ^**†**^time main effect, Tukey’s test, interval 1 > intervals 2-6, *p*s < 0.05. (**G**) Representative 20x photomicrographs of the dCA3 stratum lucidum (SL) and stratum pyramidale (SP) of rats in the Light-OFF and Light-ON groups. Brain tissue was collected immediately after the test in the cocaine-paired context. Images showing DAPI staining (blue) were used to visualize SL and SP boundaries (see also Fig. S1), and corresponding overlay images were used to visualize eYFP expression (green) and c-Fos immunoreactive (IR) cell bodies (red, *arrows*). (**H**) The density of c-Fos-immunoreactive (IR) neurons was quantified in the area corresponding to the blue rectangle on the brain schematic. c-Fos-IR cell body density (mean ± SEM) in the SL and SP of rats in the Light-ON and Light-OFF groups.

Light-ON treatment without eNpHR3.0 reduced inactive-lever responding relative to Light-OFF treatment independent of testing context (**Fig. 4E**; ANOVA, treatment main *F*_(1,14)_ = 5.62, *p* = 0.03; context main and treatment x context interaction *F*s_(1,14)_ ≤ 3.39, *p*s ≥ 0.09) due to unusually high response rate in Light-OFF controls in experiment 4 (*vs* 1-3). Time-course analysis confirmed this treatment effect, and it revealed that responding decreased over time in the cocaine-paired context at test independent of treatment type (**Fig. 4F**; ANOVA treatment main *F*_(1,14)_ = 4.41, *p* = 0.05; time main *F*_(5,70)_ = 4.00, *p* < 0.01, Tukey’s test, interval 1 > intervals 2-6, *p*s < 0.05; treatment x time interaction *F*_(5,70)_ = 2.03, *p* = 0.09).

eYFP expression was more restricted to cell bodies than eNpHR3.0-eYFP expression. Further, Light-ON treatment without eNpHR3.0 expression did not alter c-Fos expression in the SP (*t*_(14)_ = 0.52, *p* = 0.61) or SL (*t*_(14)_ = 0.48, *p* = 0.64) at test, compared to Light-OFF treatment (**Fig. 4G-H**).

### Light-ON Treatment Reduces Neuronal Activation in the dCA3 during Cocaine-Memory Reconsolidation

Experiment 5 examined whether the Light-ON treatment was sufficient to reduce neuronal activation in the dCA3 SL and SP, and specifically in GAD67-IR (GABAergic inhibitory) and CaMKII-IR (excitatory pyramidal) cell populations within these cell layers during memory reconsolidation (**Fig. 5**). Thus, brain tissue was collected immediately after memory reactivation plus 1-h optogenetic treatment. Light-ON treatment administered during the first h after memory reactivation reduced c-Fos expression in the SL (**Fig. 5C-D**; *t*_(9)_ = 2.86, *p* = 0.02) and SP (**Fig. 5C-D**; *t*_(9)_ = 6.61, *p* < 0.001), relative to Light-OFF treatment. These effects reflected reductions in GAD67+c-Fos-IR cell density in the SL (**Fig. 5E-F**; *t*_(9)_ = 2.87, *p* = 0.02) and SP (**Fig. 5E-F**; *t*_(9)_ = 5.84, *p* < 0.001) and CaMKII+c-Fos-IR cell density in the SP (**Fig. 5G-H**; *t*_(9)_ = 2.8, *p* = 0.02).

**Figure 5.**
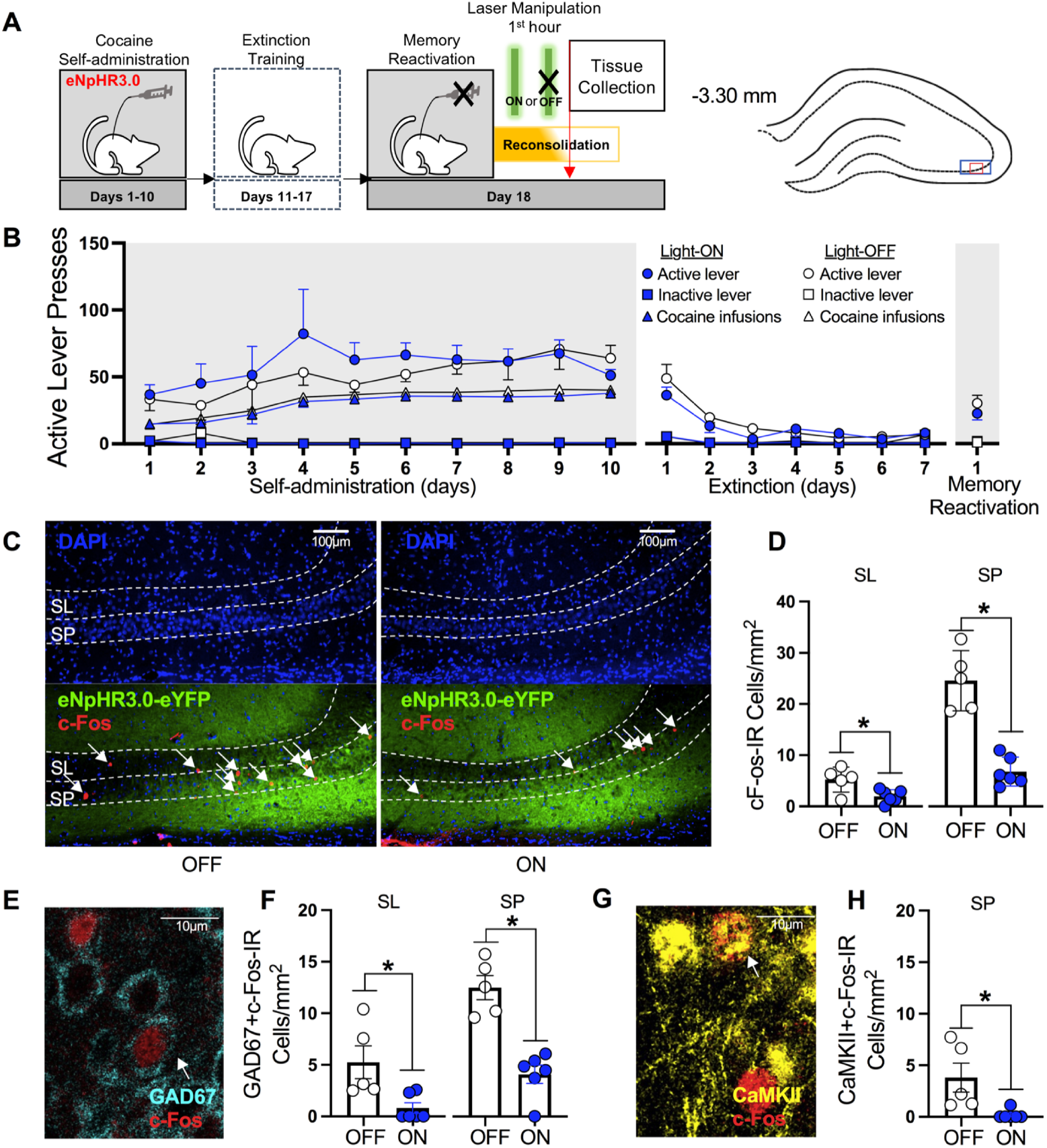
Laser-light exposure eNpHR3.0-expressing rats is sufficient to reduce neuronal activity in the dCA3. (**A**) Experimental timeline. Following cocaine self-administration and extinction training, eNpHR3.0-expressing rats received a 15-min memory-reactivation session, immediately followed by Laser-ON (n = 6) or Laser-OFF (n = 5) treatment for 1 h. Brain tissue was collected immediately after the optogenetic manipulation. The density of c-Fos-immunoreactive (IR) neurons was quantified in the areas indicated by the blue and red rectangles on the brain schematic on 20x and 40x images, respectively. (**B**) Active- and inactive-lever presses and cocaine infusions by the Light-ON and Light-OFF groups during cocaine self-administration training (last 10 days; mean/2 h ± SEM), extinction training (mean/2 h ± SEM), and during the memory-reactivation session (mean/15 min ± SEM). (**C**) Representative 20x photomicrographs of the dCA3 stratum lucidum (SL) and stratum pyramidale (SP) of rats in the Light-OFF and Light-ON groups. Images showing DAPI staining (blue) were used to visualize SL and SP boundaries (see also Fig. S1), and corresponding overlay images were used to visualize eNpHR3.0-eYFP expression (green) and c-Fos immunoreactive (IR) cell bodies (red, *arrows*). (**D**) c-Fos-IR cell body density (mean ± SEM) in the SL and SP of rats in the Light-ON and Light-OFF groups. Symbols: **\****t*-tests, *p*s < 0.05. (**E**) Representative 40x photomicrograph illustrating a GAD67+c-Fos-IR cell body (*arrow*) in the dCA3. (**F**) GAD67+c-Fos-IR cell density (mean ± SEM) in the dCA3 SL and SP of rats in the Light-ON and Light-OFF groups. Symbols: **\****t*-tests, *p*s < 0.05. (**G**) Representative 40x photomicrograph illustrating a CaMKII+c-Fos-IR cell body (*arrow*) in the dCA3. (**H**) CaMKII+c-Fos-IR cell density (mean ± SEM) in the dCA3 SP of rats in the Light-ON and Light-OFF groups. Symbol: **\****t*-test, *p* < 0.05.

## DISCUSSION

Cellular memory reconsolidation has been theorized to take place over 4-6 h after memory destabilization^26^. However, the time-dependent contributions of brain regions, such as the dCA3, to cocaine-memory reconsolidation within this time window have not been explored, likely because of the lack of spatial and temporal precision afforded by pharmacological manipulations. Here, we report that critical engagement of the dCA3 occurs during the early stages of cocaine-memory reconsolidation.

### dCA3 Plays a Requisite Role in Early-Stage Cocaine-Memory Reconsolidation

eNpHR3.0-mediated optogenetic inhibition of the dCA3 for 1 h, immediately after cocaine-memory reactivation, reduced drug context-induced cocaine-seeking behavior three days later, at test (**Fig. 1**). This decrease in cocaine-seeking behavior reflected disruption in the reconsolidation of destabilized cocaine memories, given that dCA3 inhibition without memory reactivation did not attenuate cocaine-seeking behavior (**Fig. 2**). Optogenetic inhibition of the dCA3 during the second h following memory reactivation also failed to alter cocaine-seeking behavior (**Fig. 3**), indicating that neural activity in the dCA3 is no longer required for memory reconsolidation after the first h of information processing. Diminished memory strength following optogenetic inhibition in experiment 1 did not result from heat-induced impairment in tissue health because laser-light exposure without eNpHR3.0 expression did not alter cocaine-seeking behavior (**Fig. 4**). Instead, diminished memory strength resulted from neuronal inhibition, as indicated by reduced c-Fos expression in dCA3 SL (GAD67-IR) and SP (CaMKII-IR and GAD67-IR) cell populations during memory reconsolidation (**Fig. 5**). Overall, these findings suggest that the dCA3 is recruited during the reconsolidation of labile cocaine memories in a time-limited fashion and required for the maintenance and/or subsequent recall of these memories.

Our findings expand upon literature indicating that the DH plays a critical role in memory reconsolidation. Studies have shown that pharmacological or chemogenetic inhibition of the DH impairs not only Pavlovian^27^ but also instrumental cocaine-memory reconsolidation^16^. Conversely, *de novo* protein synthesis in the DH appears to be required for memory reconsolidation in some, but not all, paradigms. For instance, inhibition of protein synthesis in the DH disrupts spatial^28^, object-recognition^29^, and morphine-conditioned place preference^30^ memory reconsolidation, but it fails to alter contextual fear-memory^31^ reconsolidation, as well as cocaine-memory reconsolidation in our instrumental model^16^. Thus, DH engagement in memory reconsolidation probably varies depending on memory age, strength, and type. In our model, the dCA3 appears to be critical during the early stages of memory reconsolidation. Thus, dCA3 neuronal activity may critically contribute to tagging post-reactivation short-term memories for restabilization^32^ or maintaining them before they can be restabilized into long-term memories.

### dCA3 SP and SL Neurons are Components of the Neural Circuitry of Cocaine-Memory Reconsolidation

Optogenetic dCA3 inhibition after cocaine-memory reactivation resulted in cell layer-specific suppression of neuronal activation in the dCA3 during memory recall, at test, depending on whether it was applied during the first or second h of memory reconsolidation (**Fig. 1-4G**). Specifically, memory impairment at test was associated with reduced SL and SP neuronal activation in experiment 1 (**Fig. 1G**), whereas intact recall was associated with no change in SL activation, with or without a decrease in SP activation, in experiments 2-4 (**Fig. 2-4G**), as indicated by c-Fos expression. Notably, the SL and SP of the dCA3 receive input primarily from dentate gyrus granule cells via mossy fibers^33,34^. Granule cells can robustly excite SP pyramidal neurons through synapses on apical dendrites proximal to cell bodies but also inhibit them indirectly, by densely innervating SL basket cells that provide feedforward inhibition onto SP pyramidal neurons^34,35^. In turn, SP pyramidal neurons excite one another, interact with SP interneurons, and stimulate dCA1 pyramidal neurons via Schaffer collaterals^33,34^. Correlated activity in these recurrent circuits during associative learning facilitates the establishment of cell ensembles that encode memory traces (i.e., engrams)^35^, and the reactivation of such engrams is required for memory reconsolidation and recall^27,36-38^. While the specific contribution of dCA3 pyramidal neurons has not been explored, studies indicate that glutamatergic pyramidal neurons in the DH are key to memory reconsolidation and subsequent recall^27.39^. For instance, CaMKII-IR pyramidal neuronal activation in the DH is necessary for Pavlovian cocaine-memory reconsolidation^26^. NR2A-containing NMDA receptor stimulation in the DH is required for cocaine-memory reconsolidation in our instrumental model^37^. Moreover, glutamatergic synaptic plasticity, specifically GluA2-containing AMPA receptor endocytosis, during reconsolidation is required for fear-memory reconsolidation^40^. The above literature and our current findings suggest that SP pyramidal neurons and SL/SP GABAergic interneurons are co-activated during, and likely interact to facilitate, cocaine-memory reconsolidation and subsequent recall. In particular, GABAergic interneurons may support these phenomena by reducing synaptic noise in associative microcircuits^35^.

Our current understanding of the larger neural circuitry within which the dCA3 supports cocaine-memory reconsolidation is limited. We have shown previously that interaction between the DH and basolateral amygdala (BLA) is necessary for cocaine-memory reconsolidation^41^. Consistent with this, unilateral inhibition of neuronal activity in the DH combined with unilateral inhibition of protein synthesis in the BLA in the contralateral hemisphere, a manipulation that functionally disconnects the DH and BLA, during reconsolidation impairs subsequent drug context-induced reinstatement and incubation of cocaine seeking^41^. Since there are no monosynaptic connections between the DH and BLA^42^, the dCA3 and BLA must interact through relay brain regions within a more complex neural circuitry. One likely relay region is the dCA1. dCA3 outputs to the dCA1 have been implicated in contextual fear-memory reconsolidation^43^, and the dCA1 and BLA exhibit synchronized theta activity during fear-memory reconsolidation^44^. Some other relay regions between the dCA1 or the dCA1-dCA3 circuit and the BLA may include the ventral hippocampus^42,45,46^, entorhinal cortex^47,48^, and perirhinal cortex^42,49^, based on their connectivity with both the BLA and DH and their recognized involvement in memory reconsolidation in other paradigms.

## CONCLUSIONS

The functional integrity of the dCA3 is required during early-stage memory reconsolidation for cocaine-memory restabilization outside of the DH. Based on extant literature^35,36^ and changes in SP and SL neuronal activation observed after the memory reactivation session and at test, we propose that cocaine-memory reconsolidation and subsequent recall are shaped by interactions between excitatory pyramidal and SL/SP inhibitory interneurons that gate the activity of associative microcircuits. To further our understanding of this process, future studies will need to examine the causal contribution of specific dCA3 cell types to cocaine-memory reconsolidation. In addition, future studies will need to systematically examine the time-dependent engagement of various brain regions during memory reconsolidation to help develop hypotheses regarding the exact contributions of those brain regions within the memory reconsolidation circuitry.

## ACKNOWLEDGMENTS

The authors are grateful for expert technical assistance provided by Justine M. C. Galliou, Peyton J. Krych, Taylor A. Brown, Jaclyn N. Roland-McGowan, Shayna R. Grogan.

## AUTHOR CONTRIBUTIONS

**SQ:** methodology, validation, investigation, data analysis, visualization, writing original draft. **SMT:** methodology, investigation, validation, data analysis, writing original draft. **RW:** investigation, validation. **JAH:** methodology, validation, investigation. **JLR:** methodology, investigation, visualization. **CKI:** methodology, investigation. **AAA:** methodology, validation, investigation. **RJC:** investigation. **RFL:** conceptualization, methodology, data analysis, writing original draft, review and editing, supervision, funding acquisition.

## REFERENCES

1. Dackis CA, O’Brien CP. Cocaine dependence: A disease of the brain’s reward centers. J Subst Ab Treat. 2001;21:111–117.

2. Hyman SE. Addiction: A disease of learning and memory. Am J Psych. 2005;162:1414–1422.

3. Fuchs RA, Lasseter HC, Ramirez DR, Xie X. Relapse to drug seeking following prolonged abstinence: the role of environmental stimuli. Drug Discov Today Dis Models. 2009;5:251–258.

4. Bender BN, Torregrossa MM. Molecular and circuit mechanisms regulating cocaine memory. Cell Mol Life Sci. 2020;0123456789.

5. Ehrman RN, Robbins SJ, Childress AR, O’Brien CP. Conditioned responses to cocaine-related stimuli in cocaine abuse patients. Psychopharmacology (Berl). 1992;107:523–529.

6. Tronson NC, Taylor JR. Molecular mechanisms of memory reconsolidation. Nat Rev Neurosci. 2007;8:262–275.

7. Finnie PSB, Nader K. The role of metaplasticity mechanisms in regulating memory destabilization and reconsolidation. Neurosci Biobeh Rev. 2012;36:1667–1707.

8. Saladin ME, Gray KM, McRae-Clark AL, Larowe SD, Yeatts SD., Baker NL, et al. A double blind, placebo-controlled study of the effects of post-retrieval propranolol on reconsolidation of memory for craving and cue reactivity in cocaine dependent humans. Psychopharmacology (Berl). 2014;226:721–737.

9. Lonergan M, Saumier D, Tremblay J, Kieffer B, Brown TG, Brunet A. Reactivating addiction-related memories under propranolol to reduce craving: A pilot randomized controlled trial. J Beh Therapy Exp Psych. 2016;50:245–249.

10. Taylor J R, Olausson P, Quinn JJ, Torregrossa MM. Targeting extinction and reconsolidation mechanisms to combat the impact of drug cues on addiction. Neuropharmacology. 2009;56(Suppl. 1):186–195.

11. Sorg BA. Reconsolidation of drug memories. Neurosci Biobeh Rev. 2012;36:1400–1417.

12. Exton-McGuinness MTJ, Milton AL. Reconsolidation blockade for the treatment of addiction: Challenges, new targets, and opportunities. Learn Mem. 2018;25:492–500.

13. Lee JLC, Everitt BJ, Thomas KL. Independent cellular processes for hippocampal memory consolidation and reconsolidation. Science. 2004;304:839–843.

14. Solstad T, Moser EI, Einevoll GT. Inhibition of mRNA synthesis in the hippocampus impairs consolidation and reconsolidation of spatial memory. Hippocampus. 2006;1031:1026–1031.

15. Lee JLC. Memory reconsolidation mediates the strengthening of memories by additional learning. Nat Neurosci. 2008;11:1264–1266.

16. Ramirez DR, Bell GH, Lasseter HC, Xie X., Traina SA, Fuchs RA. Dorsal hippocampal regulation of memory reconsolidation processes that facilitate drug context-induced cocaine-seeking behavior in rats. Eur J Neurosci. 2009;30:1–7.

17. Wells AM, Xie X, Higginbotham JA, Arguello AA, Healey KL, Blanton ^M^, Fuchs RA. Contribution of an SFK-mediated signaling pathway in the dorsal hippocampus to cocaine-memory reconsolidation in rats. Neuropsychopharmacology. 2016;41:675–85.

18. Kesner RP. A process analysis of the CA3 subregion of the hippocampus. Front Cell Neurosci. 2013;7:1–17.

19. Le Duigou C, Simonnet J, Teleñczuk MT, Fricker D, Miles R. Recurrent synapses and circuits in the CA3 region of the hippocampus: An associative network. Front Cell Neurosci. 2014;7:1–13.

20. Lines J, Nation K, Fellous JM. Dorsoventral and proximodistal hippocampal processing account for the influences of sleep and context on memory (re)consolidation: A connectionist model. Comput Intell Neurosci. 2017; 2017:8091780.

21. Huang B, Zhu H, Zhou Y, Liu X, Ma L. Unconditioned- and conditioned-stimuli induce differential memory reconsolidation and β-AR-dependent CREB activation. Front Neural Circuits. 2017;11:1– 10.

22. Fuchs RA, Evans KA, Ledford CC, Parker MP, Case JM, Mehta RH, See RE. The role of the dorsomedial prefrontal cortex, basolateral amygdala, and dorsal hippocampus in contextual reinstatement of cocaine seeking in rats. Neuropsychopharmacology. 2005;30:296–309.

23. Bott, JB, Héraud C, Cosquer B, Herbeaux K, Aubert J, Sartor M, et al. APOE-sensitive cholinergic sprouting compensates for hippocampal dysfunctions due to reduced entorhinal input. J Neurosci. 2016;36:10472–10486

24. Dȩbiec J, Doyère V, Nader K, LeDoux JE. Directly reactivated, but not indirectly reactivated, memories undergo reconsolidation in the amygdala. Proc Natl Acad Sci U S A. 2006;103:3428– 3433.

25. Tye KM, Deisseroth K. Optogenetic investigation of neural circuits underlying brain disease in animal models. Nat Rev Neurosci. 2012;13:251–266.

26. Nader K, Hardt O. A single standard for memory: The case for reconsolidation. Nat Rev Neurosci. 2009;10:224–234.

27. Liu C, Sun X, Wang Z, Le Q, Liu P, Jiang C, et al. Retrieval-induced upregulation of Tet3 in pyramidal neurons of the dorsal hippocampus mediates cocaine-associated memory reconsolidation. Int J Neuropsychopharmacol. 2018;21:255–266.

28. Rossato JI, Bevilaqua LRM, Medina JH, Izquierdo I, Cammarota M. Retrieval induces hippocampal-dependent reconsolidation of spatial memory. Learn Mem. 2006;13:431–440.

29. Rossato JI, Bevilaqua LR, Myskiw JC, Medina JH, Izquierdo I, Cammarota M. On the role of hippocampal protein synthesis in the consolidation and reconsolidation of object recognition memory. Learn Mem. 2007;14:36–46.

30. Milekic MH, Brown SD, Castellini C, Alberini CM. Persistent disruption of an established morphine conditioned place preference. J Neurosci. 2006;26:3010–3020.

31. Biedenkapp JC, Rudy JW. Context memories and reactivation: Constraints on the reconsolidation hypothesis. Behav Neurosci. 2004;118:956–964.

32. Rabinovich Orlandi I, Fullio CL, Schroeder MN, Giurfa M, Ballarini F, Moncada D. Behavioral tagging underlies memory reconsolidation. PNAS. 2020;117:18029–18036.

33. Spruston N, Lübke J, Frotscher M. Interneurons in the stratum lucidum of the rat hippocampus: An anatomical and electrophysiological characterization. J Comp Neurol.1997;385:427–440.

34. Le Duigou C, Simonnet J, Teleñczuk MT, Fricker D, Miles R. Recurrent synapses and circuits in the CA3 region of the hippocampus: an associative network. Front Cell Neurosci. 2014;7:262.

35. Apóstolo N, Smukowski SN, Vanderlinden J, Condomitti G, Rybakin V, Ten Bos J. Synapse type-specific proteomic dissection identifies IgSF8 as a hippocampal CA3 microcircuit organizer. Nat Comm. 2020;11:5171.

36. Kim J, Kwon JT, Kim HS, Josselyn SA, Han JH. Memory recall and modifications by activating neurons with elevated CREB. Nat Neurosci. 2014;17:65–72.

37. Sekeres MJ, Mercaldo V, Richards B, Sargin D, Mahadevan V, Woodin MA, Frankland PW, Josselyn SA. Increasing CRTC1 function in the dentate gyrus during memory formation or reactivation increases memory strength without compromising memory quality. J Neurosci. 2012;32:17857–68.

38. Yousuf M, Packard PA, Fuentemilla L, Bunzeck N. Functional coupling between CA3 and laterobasal amygdala supports schema dependent memory formation. NeuroImage. 2021;244:118563.

39. Wells, AM, Xie X, Higginbotham JA, Arguello AA, Healey KL, Blanton M, Fuchs RA. Contribution of an SFK-mediated signaling pathway in the dorsal hippocampus to cocaine-memory reconsolidation in rats. Neuropsychopharmacology. 2016;41:675–685.

40. Rao-Ruiz P, Rotaru DC, Van Der Loo RJ, Mansvelder HD, Stiedl O, Smit AB, Spijker S. Retrieval-specific endocytosis of GluA2-AMPARs underlies adaptive reconsolidation of contextual fear. Nat Neurosci. 2011;14:1302–1308.

41. Wells AM, Lasseter HC, Xie X, Cowhey KE, Reittinger AM, Fuchs RA. Interaction between the basolateral amygdala and dorsal hippocampus is critical for cocaine memory reconsolidation and subsequent drug context-induced cocaine-seeking behavior in rats. Learn Mem. 2011;18:693– 702.

42. Pitkänen A, Pikkarainen M, Nurminen N, Ylinen A. Reciprocal connections between the amygdala and the hippocampal formation, perirhinal cortex, and postrhinal cortex in rat. Ann N Y Acad Sci. 2000;911:369–391.

43. Lux V, Masseck OA, Herlitze S, Sauvage MM. Optogenetic destabilization of the memory trace in CA1: Insights into reconsolidation and retrieval processes. Cereb Cortex. 2017;27:841–851.

44. Narayanan RT, Seidenbecher T, Sangha S, Stork O, Pape HC. Theta resynchronization during reconsolidation of remote contextual fear memory. NeuroReport. 2007;18:1107–1111.

45. Bienkowski MS, Bowman I, Song MY, Gou L, Ard T, Cotter K, et al. Integration of gene expression and brain-wide connectivity reveals the multiscale organization of mouse hippocampal networks. Nat Neurosci. 2018;21:1628–1643.

46. Chia C, Otto T. Hippocampal Arc (Arg3.1) expression is induced by memory recall and required for memory reconsolidation in trace fear conditioning. Neurobiol Learn Mem. 2013;106:48–55.

47. Baldi E, Bucherelli C. Entorhinal cortex contribution to contextual fear conditioning extinction and reconsolidation in rats. Neurobiol Learn Mem. 2014;110:64–71.

48. Wahlstrom KL, Alvarez-Dieppa AC, McIntyre CK, LaLumiere RT. The medial entorhinal cortex mediates basolateral amygdala effects on spatial memory and downstream activity-regulated cytoskeletal-associated protein expression. Neuropsychopharmacology. 2021;46:1172–1182.

49. Winters BD, Tucci MC, Jacklin DL, Reid JM, Newsome J. On the dynamic nature of the engram: Evidence for circuit-level reorganization of object memory traces following reactivation. J Neurosci. 2011;31:17719–17728.

