## Supplemental Methods and Figure S1 for "Optogenetic inhibition of the dorsal hippocampus CA3 region during early-stage cocaine-memory reconsolidation disrupts subsequent context-induced cocaine seeking in rats"

### Supplemental Methods

#### *Food Training*

Food training was undertaken to expedite the acquisition of lever responding for un-signaled intravenous cocaine infusions at a later stage of the experiment. Rats were acclimated to handling for at least three consecutive days prior to food training. Food training occurred in operant-conditioning chambers (26 × 27 × 27 cm; Coulbourn Instruments, Allentown, PA) equipped with two levers. During the food-training session, each response on a designated lever (active lever) resulted in food reinforcement (45 mg pellets; Bio-Serv, Flemington, NJ) under a continuous reinforcement schedule. Presses on a second lever (inactive lever) did not result in food reinforcement. Training continued until the rats obtained at least 100 food pellets during a session (mean ± SEM = 1.38 ± 0.63 sessions). During food training, the rats had no access to the contextual stimuli that were used during subsequent cocaine self-administration training.

#### *Virus Microinfusion Procedure*

Virus microinfusions were delivered through 5-μl Hamilton syringes fitted with 26-gauge needles, using a nanoinfusion pump mounted on the stereotaxic instrument. The needles remained at the injection site for 2 min before and for 10 minutes after each infusion.

#### *Intravenous Catheter Surgery*

Up to three days after the stereotaxic surgery, rats were anesthetized using ketamine and xylazine (100 mg/kg and 5 mg/kg, i.p., respectively). Catheters were constructed in-house and implanted into the right jugular vein as described previously[22]. The catheter tubing ran under the skin and exited posterior to the rat's shoulder blades. The rats received bacon-flavored Rimadyl® MD tablets for at least 48 h after the catheter surgery. The rats were given at least five days for post-operative recovery. The catheters were flushed daily with 0.1 ml of a cefazolin antibiotic solution (10.0 mg/ml), followed by 0.1 ml of heparinized saline (70 U/ml) to maintain catheter patency. Catheter patency was tested using propofol (1 mg/0.1 ml), which induces rapid loss of muscle tone when administered intravenously.

#### ***Training Contexts***

The operant conditioning chambers were modified to construct two distinct environmental contexts in separate behavioral testing rooms. The two contexts were used for drug self-administration and extinction training in a counterbalanced manner across rats in each treatment group. Context 1 contained a continuous red light, intermittent tone (78 dB, 10 Hz, 2 sec on, 2 sec off), a vanilla-scented air freshener, and wire mesh flooring (26 cm x 27 cm). Context 2 contained an intermittent white light stimulus above the inactive lever, a continuous tone stimulus (78 dB, 2kHz, 2 sec on, 4 sec off), a pine-scented air freshener, and steel grid flooring bisected by a slanted ceramic tile (19 cm x 27 cm).

#### ***Acclimation Procedure***

Immediately after extinction sessions 5, 6, and 7, rats were acclimated to the optogenetic procedure. To this end, rats were placed into novel, black metal chambers (15 x 11 x 20"). Their indwelling optic fibers were connected to fiber optic patch cables (Thor Labs). The other end of the patch cables was connected to an optical communicator (Doric Lenses, Quebec, CAN) that was suspended above the chamber. The rats remained in the chambers for 1 h without laser-light stimulation.

#### *Immunohistochemistry*

In experiments 1-4, c-Fos expression was assessed in the dCA3 in tissue collected at test. Free-floating brain sections were first blocked in 5% normal donkey and goat serum in 0.5% phosphate-buffered saline (PBS) with 3% Triton X for 30 min. The sections were then incubated with mouse anti-c-Fos (1:500; sc-271243, Santa Cruz Biotechnology, Dallas, TX) and chicken anti-GFP (1:500; GFP697986, Aves Labs, Tigard, OR) primary antibodies at room temperature (RT) for 24 h. Next, the sections were washed in PBS and then incubated with a secondary antibody solution containing donkey anti-mouse-cy3 (1:500; 715-165-150, Jackson ImmunoResearch, West Grove, PA) and goat anti-chicken-488 (1:1000; A11039, ThermoFisher Scientific, Waltham, MA) secondary antibodies at RT for 2 h. AT last, the sections were stained with DAPI for 5 minutes (1:1000; DAPI solution, 62248, ThermoFisher Scientific). Brain sections were mounted on glass slides, preserved with ProLong Diamond antifade mountant (ThermoFisher Scientific, Waltham, MA), and coverslipped. Three images of the dCA3, ventral to the tip of the optic fiber, were obtained using a 10x objective for each rat (Leica DMI6000 B, Leica, Wetzlar, Germany). Uniform background subtraction and contrast enhancement were applied in ImageJ. c-Fos-IR cell body density (cell bodies/mm$^2$) was quantified ventral to the optic fiber tract, within the SL and SP cell layers of the dCA3, using a custom macro in ImageJ. Cell density values were averaged across the three images.

In experiment 5, c-Fos expression was assessed in tissue collected after memory reactivation and the 1-h optogenetic manipulation (i.e., during cocaine-memory reconsolidation), and c-Fos expression was characterized in the SL and SP, and specifically in CaMKII-IR cell populations in the SP and in GAD67-IR cell populations in the SP and SL. Brain sections were blocked in 5% normal donkey and goat serum in 0.5% phosphate-buffered saline (PBS) with 3% Triton X for 30 min. Separate sets of brain sections were then incubated with mouse anti-GAD67 (1:1000; MAB5406, Sigma-Aldrich, St. Louis, MO) or mouse anti-CaMKII (1:200; sc5306, Santa Cruz Biotechnology) primary antibodies at RT for 24 h. Next, the brain sections were incubated with AffiniPure fab fragment rabbit-anti-mouse IgG (1:100; 315-007-003, Jackson ImmunoResearch, West Grove, PA) at RT for 30 minutes to prevent cross-binding during c-Fos staining. Brain sections were then incubated with donkey anti-rabbit-680 (1:500; 926-68023, Li-Cor, Lincoln, NE) secondary antibody at RT for 2 h. Next, both sets of brain sections were incubated with mouse anti-c-Fos (1:500; sc-271243, Santa Cruz Biotechnology, Dallas, TX) primary antibody then donkey anti-mouse-cy3 (1:500; 715-165-150, Jackson ImmunoResearch) secondary antibody. Brain sections were mounted on glass slides, preserved with ProLong Diamond antifade mountant (ThermoFisher Scientific), and coverslipped. Two confocal z-stack images were obtained using a 20x objective (14 steps, z-step size 0.35 μm; Leica SP8; Leica, Wetzlar, Germany) for each brain section. After uniform background subtraction and contrast enhancement, GAD67+c-Fos-IR and CaMKII+c-Fos-IR cell body density (cell bodies/mm$^2$) was visualized in the dCA3, ventral to the optic fiber tract. C-Fos-IR cell bodies were quantified, and cell density values were averaged over two z-stack max projection images using a custom macro in ImageJ. Double-labeled cells were hand counted in each step, and density values were averaged over the two z-stack images.

Figure S1

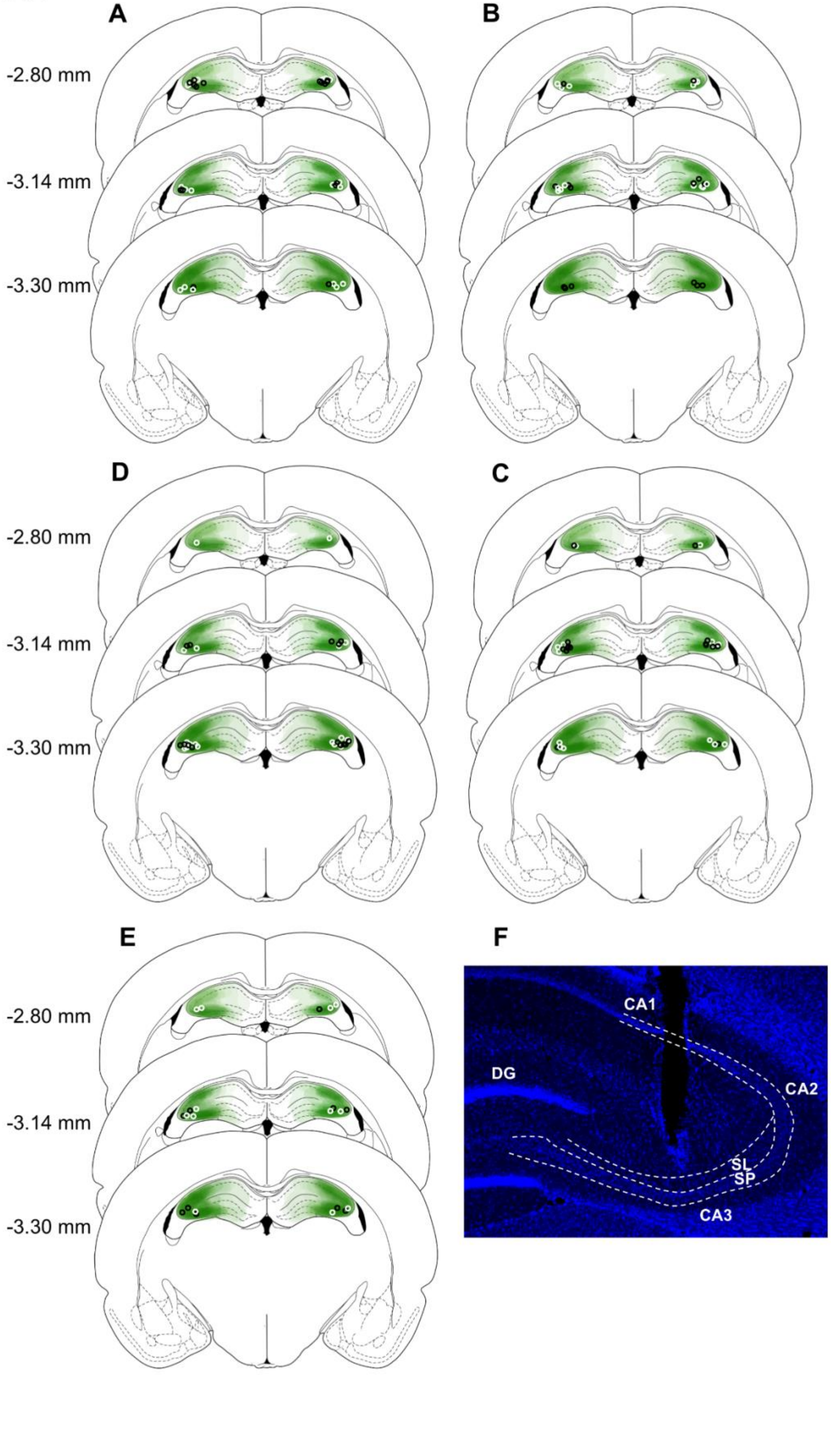
A
B
-2.80 mm
-3.14 mm
-3.30 mm
D
C
-2.80 mm
-3.14 mm
-3.30 mm
E
F
-2.80 mm
-3.14 mm
-3.30 mm
CA1
DG
CA2
SL
SP
CA3

**Figure S1.** Schematic illustration of eNpHR3.0-eYFP expression in experiments 1-3 and 5 (green shading in **A**, **B**, **C** and **E**, respectively), eYFP expression in experiment 4 (green shading in **D**), and the most ventral point of optic fiber-tracts for all rats in the Light-OFF (*black symbols*) and Light-ON (*white symbols*) groups in each experiment. The numbers next to the schematics represent the AP distance relative to bregma. (**F**) Representative 5x photomicrograph of a DAPI-stained DH section showing an optic-fiber track in relation to the stratum lucidum (SL) and stratum pyramidale (SP) cell layers of the dCA3. SP and SL boundaries were identified based on differences in cell density[23].

**Table S1. Behavioral History in Experiments 1-5**

| **Experiment 1** | | | | | | | | | | | | | |
|---|---|---|---|---|---|---|---|---|---|---|---|---|---|
| **Phase** | **Measure** | **Treatment Main Effects** | | | | **Time Main Effects** | | | | **Treatment x Time** | | | |
| | | **Test** | ***df*** | **Statistic** | ***p*** | **Test** | ***df*** | **Statistic** | ***p*** | **Test** | ***df*** | **Statistic** | ***p*** |
| **Self-Administration[a]** | Active Lever | *F* | 1,15 | 3.69 | 0.07 | *F* | 9,135 | 0.45 | 0.90 | *F* | 9,135 | 1.56 | 0.13 |
| | Inactive Lever | *F* | 1,15 | 0.14 | 0.71 | *F* | 9,135 | 1.79 | 0.10 | *F* | 9,135 | 2.67 | 0.01 |
| | Cocaine Infusions | *F* | 1,15 | 3.12 | 0.10 | *F* | 9,135 | 18.84 | <0.001 | *F* | 9,135 | 1.89 | 0.06 |
| **Extinction[b]** | Active Lever | *F* | 1,15 | 1.46 | 0.25 | *F* | 6,90 | 35.39 | <0.001 | *F* | 6,90 | 2.97 | 0.01 |
| | Inactive Lever | *F* | 1,15 | 0.18 | 0.68 | *F* | 6,90 | 7.03 | <0.001 | *F* | 6,90 | 0.40 | 0.90 |
| **Memory Reactivation[c]** | Active Lever | *t* | 15 | -0.20 | 0.85 | | | | | | | | |
| | Inactive Lever | *t* | 15 | 1.22 | 0.24 | | | | | | | | |
| **Experiment 2** | | | | | | | | | | | | | |
| **Phase** | **Measure** | **Treatment Main Effects** | | | | **Time Main Effects** | | | | **Treatment x Time** | | | |
| | | **Test** | ***df*** | **Statistic** | ***p*** | **Test** | ***df*** | **Statistic** | ***p*** | **Test** | ***df*** | **Statistic** | ***p*** |
| **Self-Administration[a]** | Active Lever | *F* | 1,12 | 1.03 | 0.33 | *F* | 9,108 | 5.33 | <0.001 | *F* | 9,108 | 2.21 | 0.03 |
| | Inactive Lever | *F* | 1,12 | 0.11 | 0.75 | *F* | 9,108 | 1.15 | 0.34 | *F* | 9,108 | 0.86 | 0.56 |
| | Cocaine Infusions | *F* | 1,12 | 9.54 | 0.01 | *F* | 9,108 | 51.51 | <0.001 | *F* | 9,108 | 1.28 | 0.26 |

| Phase | Measure | Treatment Main Effects | | | | Time Main Effects | | | | Treatment x Time | | | |
|---|---|---|---|---|---|---|---|---|---|---|---|---|---|
| Extinction[b] | Active Lever | *F* | 1,12 | 0.80 | 0.39 | *F* | 6,72 | 17.94 | <0.001 | *F* | 6,72 | 0.82 | 0.56 |
| | Inactive Lever | *F* | 1,12 | 0.27 | 0.61 | *F* | 6,72 | 3.84 | <0.001 | *F* | 6,72 | 0.32 | 0.93 |
| No-Memory Reactivation[c] | Active Lever | | | | | | | | | | | | |
| | Inactive Lever | | | | | | | | | | | | |
| **Experiment 3** | | | | | | | | | | | | | |
| **Phase** | **Measure** | **Treatment Main Effects** | | | | **Time Main Effects** | | | | **Treatment x Time** | | | |
| | | **Test** | ***df*** | **Statistic** | ***p*** | **Test** | ***df*** | **Statistic** | ***p*** | **Test** | ***df*** | **Statistic** | ***p*** |
| **Self-Administration[a]** | Active Lever | *F* | 1,14 | 0.21 | 0.65 | *F* | 9,126 | 0.54 | 0.84 | *F* | 9,126 | 0.73 | 0.68 |
| | Inactive Lever | *F* | 1,14 | 0.46 | 0.51 | *F* | 9,126 | 1.81 | 0.07 | *F* | 9,126 | 0.90 | 0.53 |
| | Cocaine Infusions | *F* | 1,14 | 2.15 | 0.17 | *F* | 9,126 | 9.39 | <0.001 | *F* | 9,126 | 1.23 | 0.28 |
| **Extinction[b]** | Active Lever | *F* | 1,14 | 0.74 | 0.40 | *F* | 6,84 | 9.94 | <0.001 | *F* | 6,84 | 0.27 | 0.96 |
| | Inactive Lever | *F* | 1,14 | 10.00 | 0.01 | *F* | 6,84 | 2.45 | 0.03 | *F* | 6,84 | 0.59 | 0.74 |
| **Memory Reactivation[c]** | Active Lever | *t* | 14 | 0.41 | 0.69 | | | | | | | | |
| | Inactive Lever | *t* | 14 | -1.10 | 0.31 | | | | | | | | |
| **Experiment 4** | | | | | | | | | | | | | |
| **Phase** | **Measure** | **Treatment Main Effects** | | | | **Time Main Effects** | | | | **Treatment x Time** | | | |
| | | **Test** | ***df*** | **Statistic** | ***p*** | **Test** | ***df*** | **Statistic** | ***p*** | **Test** | ***df*** | **Statistic** | ***p*** |
| **Self-Administration[a]** | Active Lever | *F* | 1,14 | 0.10 | 0.76 | *F* | 9,126 | 1.28 | 0.26 | *F* | 9,126 | 0.77 | 0.65 |
| | Inactive Lever | *F* | 1,14 | 0.18 | 0.68 | *F* | 9,126 | 1.16 | 0.33 | *F* | 9,126 | 0.79 | 0.63 |
| | Cocaine Infusions | *F* | 1,14 | 0.52 | 0.48 | *F* | 9,126 | 8.50 | <0.001 | *F* | 9,126 | 0.38 | 0.94 |
| **Extinction[b]** | Active Lever | *F* | 1,14 | 0.02 | 0.88 | *F* | 6,84 | 28.14 | <0.001 | *F* | 6,84 | 1.53 | 0.18 |
| | Inactive Lever | *F* | 1,14 | 3.68 | 0.08 | *F* | 6,84 | 9.03 | <0.001 | *F* | 6,84 | 0.88 | 0.51 |
| **Memory Reactivation[c]** | Active Lever | *t* | 14 | 0.30 | 0.77 | | | | | | | | |
| | Inactive Lever | *t* | 14 | 0.11 | 0.91 | | | | | | | | |
| **Experiment 5** | | | | | | | | | | | | | |

| Phase | Measure | Treatment Main Effects | | | | Time Main Effects | | | | Treatment x Time | | | |
|---|---|---|---|---|---|---|---|---|---|---|---|---|---|
| | | **Test** | ***df*** | **Statistic** | ***p*** | **Test** | ***df*** | **Statistic** | ***p*** | **Test** | ***df*** | **Statistic** | ***p*** |
| **Self-Administration[a]** | Active Lever | *F* | 1, 9 | 0.41 | 0.54 | *F* | 9,81 | 1.97 | 0.05 | *F* | 9,81 | 0.52 | 0.85 |
| | Inactive Lever | *F* | 1,9 | 0.86 | 0.38 | *F* | 9,81 | 1.47 | 0.17 | *F* | 9,81 | 1.32 | 0.24 |
| | Cocaine Infusions | *F* | 1.9 | 1.48 | 0.26 | *F* | 9,81 | 12.80 | <0.001 | *F* | 9,81 | 0.14 | 1.00 |
| **Extinction[b]** | Active Lever | *F* | 1,9 | 1.35 | 0.28 | *F* | 6,54 | 20.13 | <0.001 | *F* | 6.54 | 1.03 | 0.41 |
| | Inactive Lever | *F* | 1,9 | 0.97 | 0.35 | *F* | 6.54 | 3.59 | 0.01 | *F* | 6.54 | 1.35 | 0.10 |
| **Memory Reactivation[c]** | Active Lever | *t* | 9 | -1.00 | 0.35 | | | | | | | | |
| | Inactive Lever | *t* | 9 | 0.41 | 0.69 | | | | | | | | |

[a]Lever responses and cocaine infusions across the last 10 self-administration training days were analyzed using 2 x 10 mixed-factorial ANOVAs. [b]Lever responses across the 7 extinction training days prior to memory reactivation were analyzed using 2 x 7 mixed-factorial ANOVAs. [c]The total number of lever responses during the 15-min memory-reactivation session was analyzed using independent samples *t*-tests.
